## Supplementary figures and images for "Establishment of quantitative RNAi-based forward genetics in *Entamoeba histolytica* and identification of genes required for growth"

### Figure S1

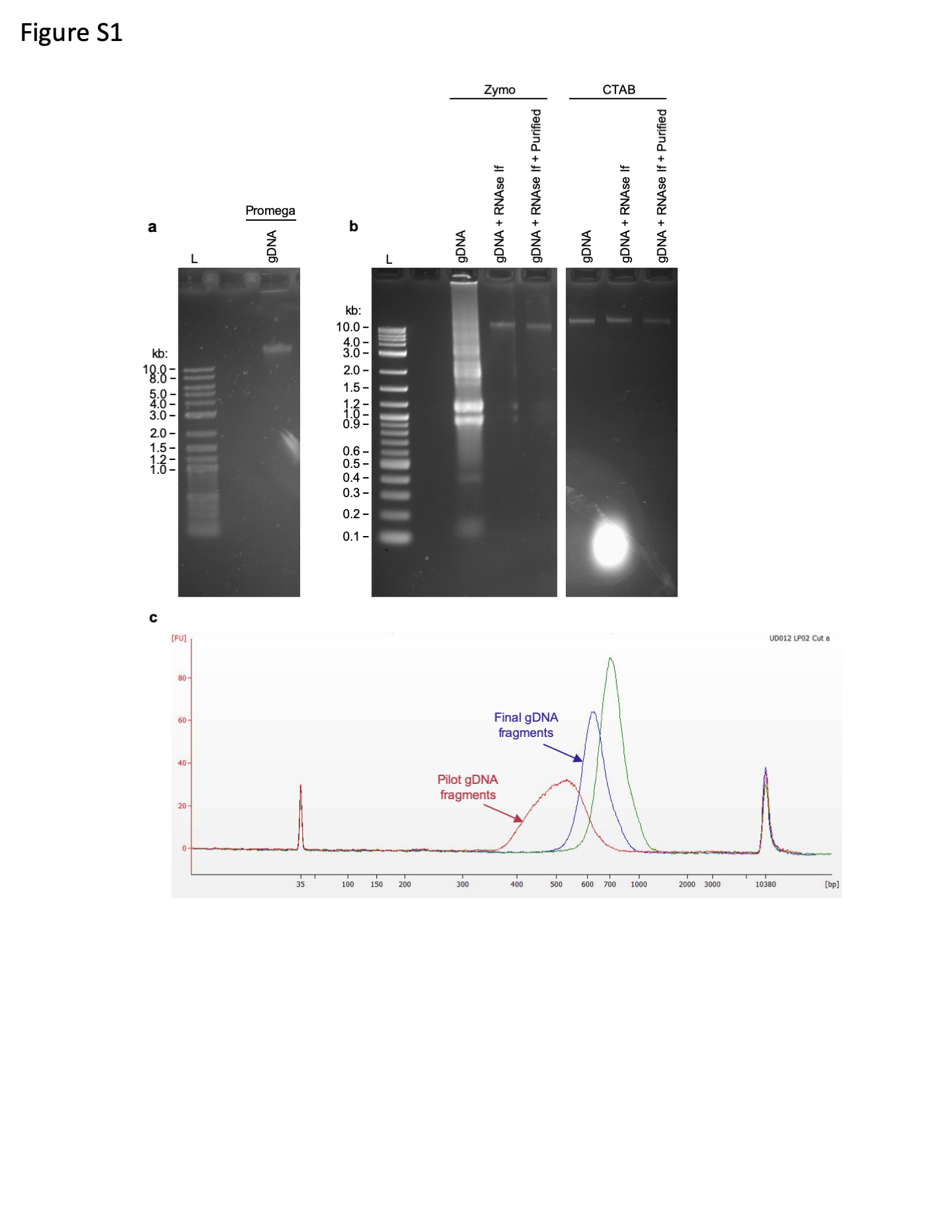

### Figure S2

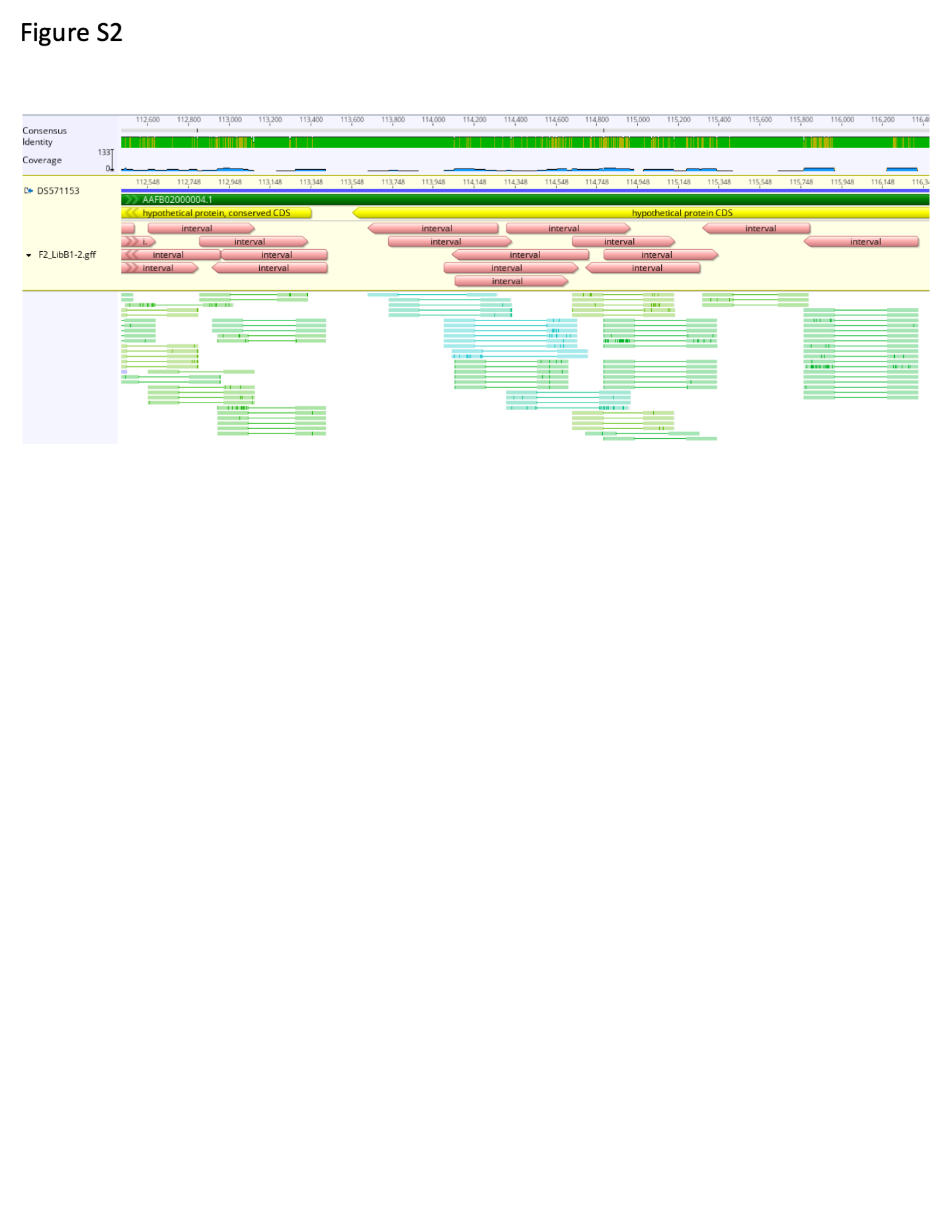

### Figure S3

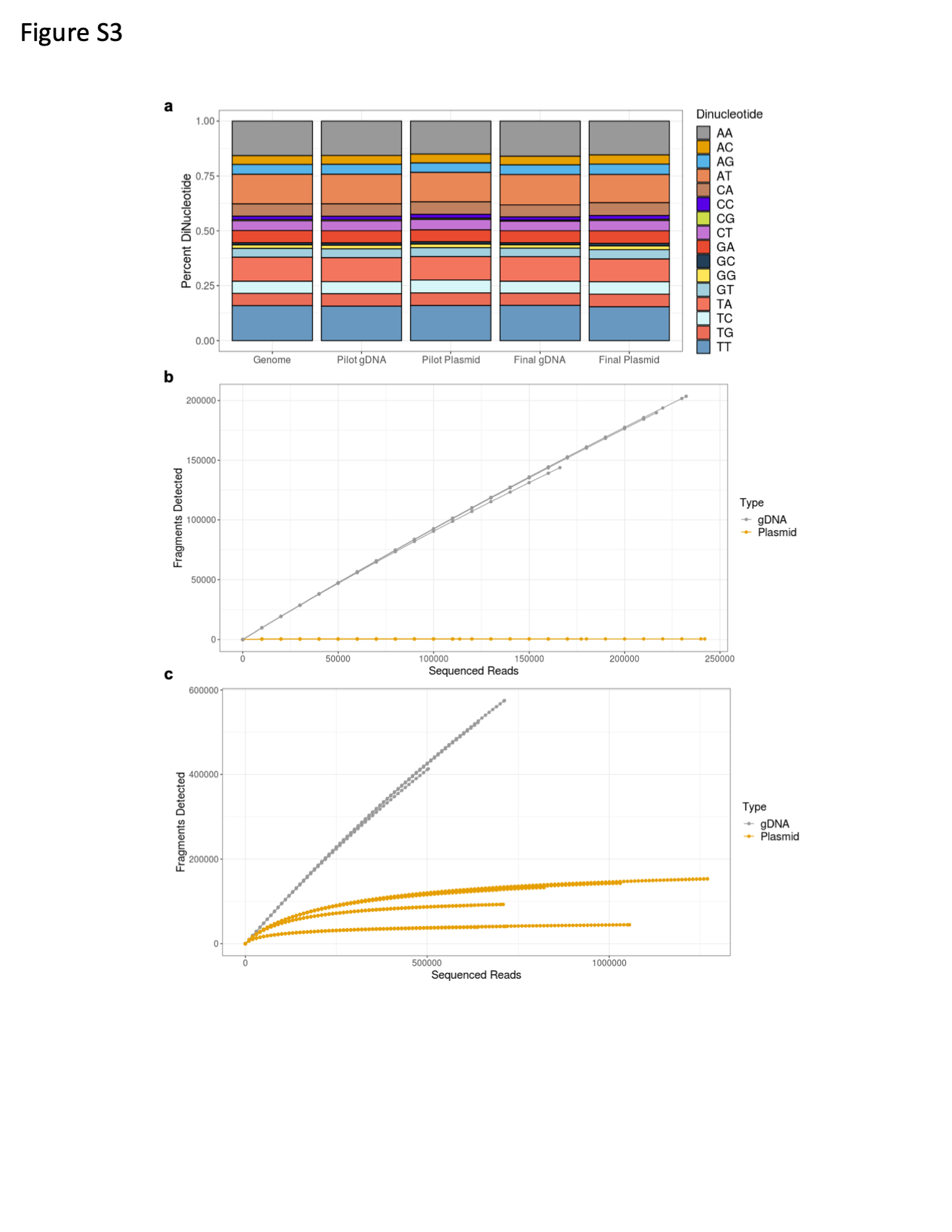

### Figure S4

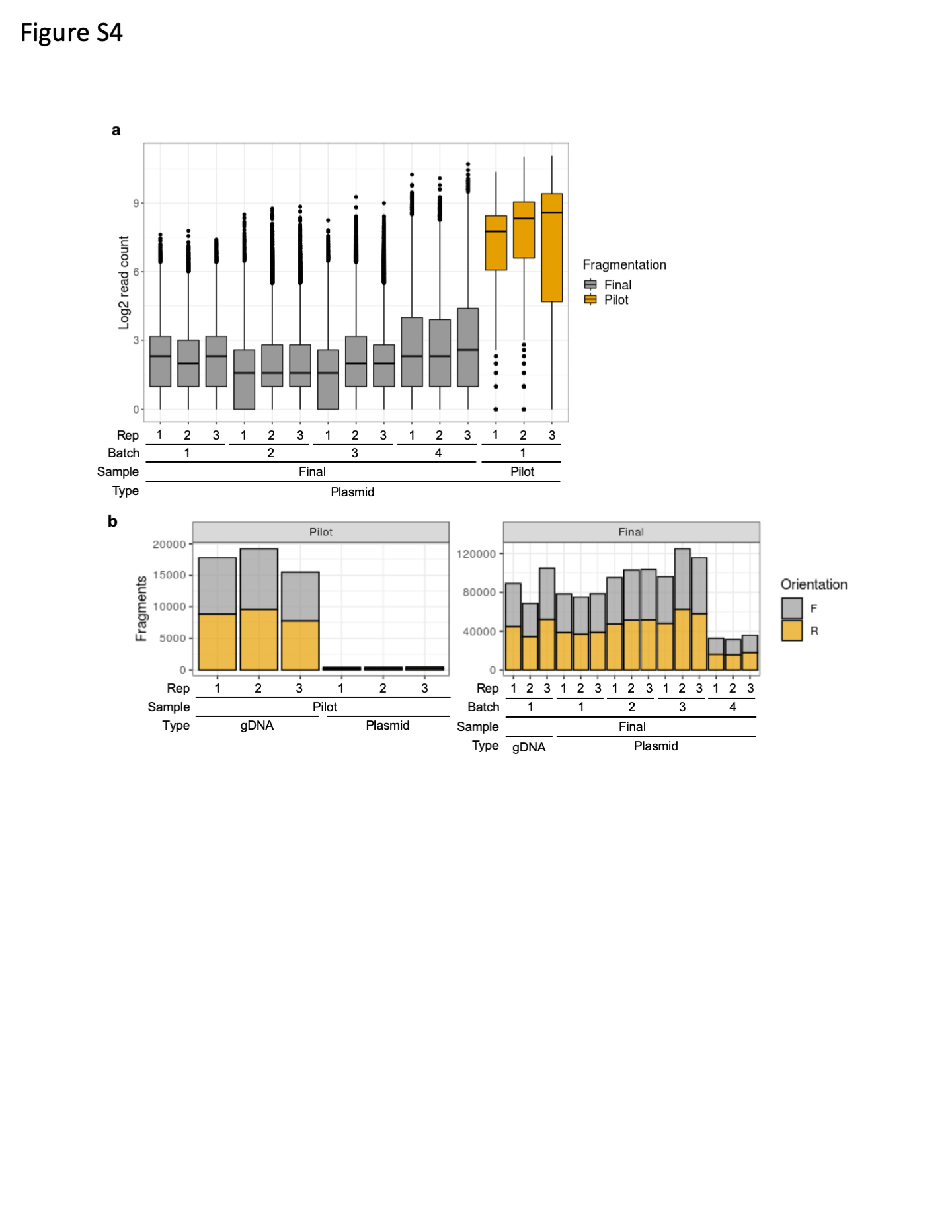

### Figure S5

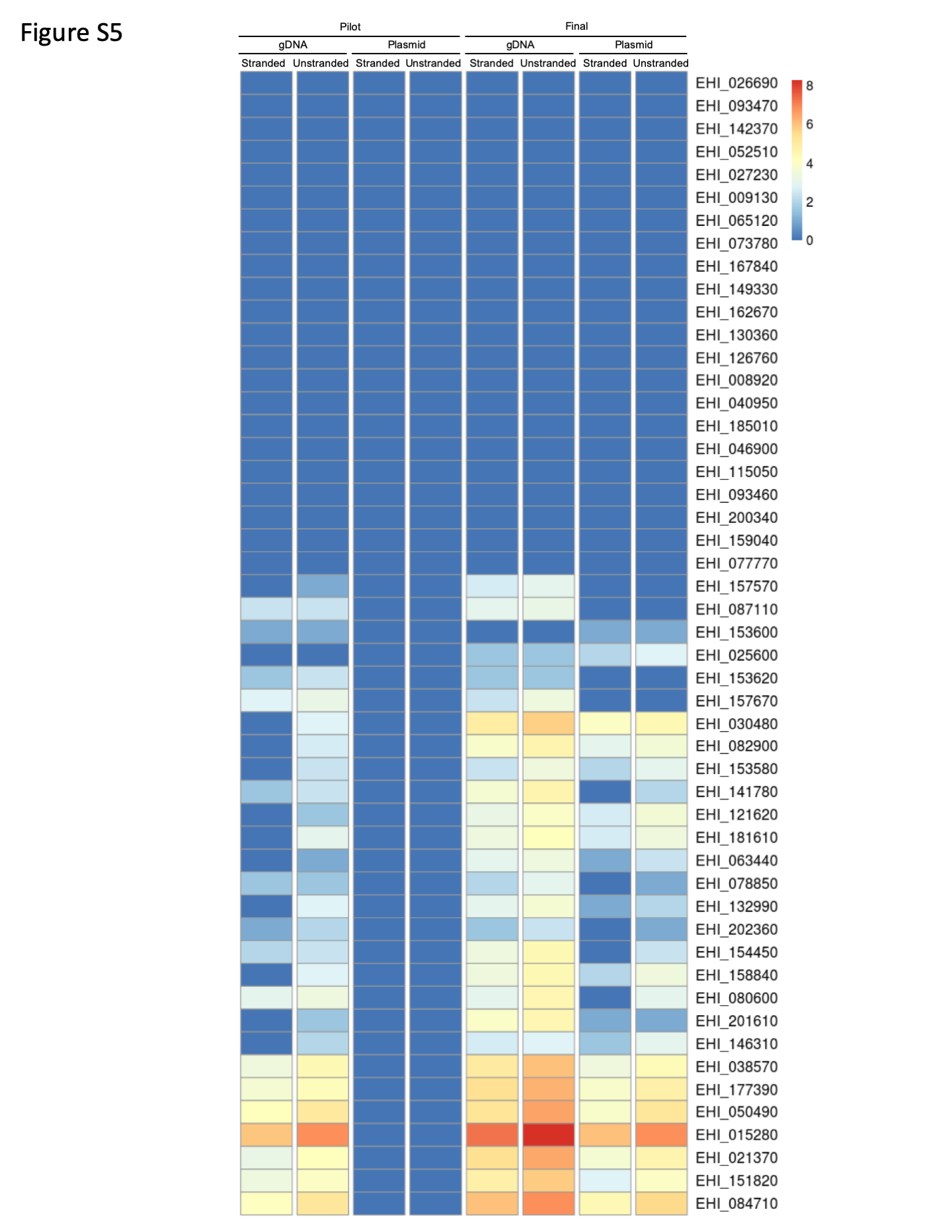

### Figure S6

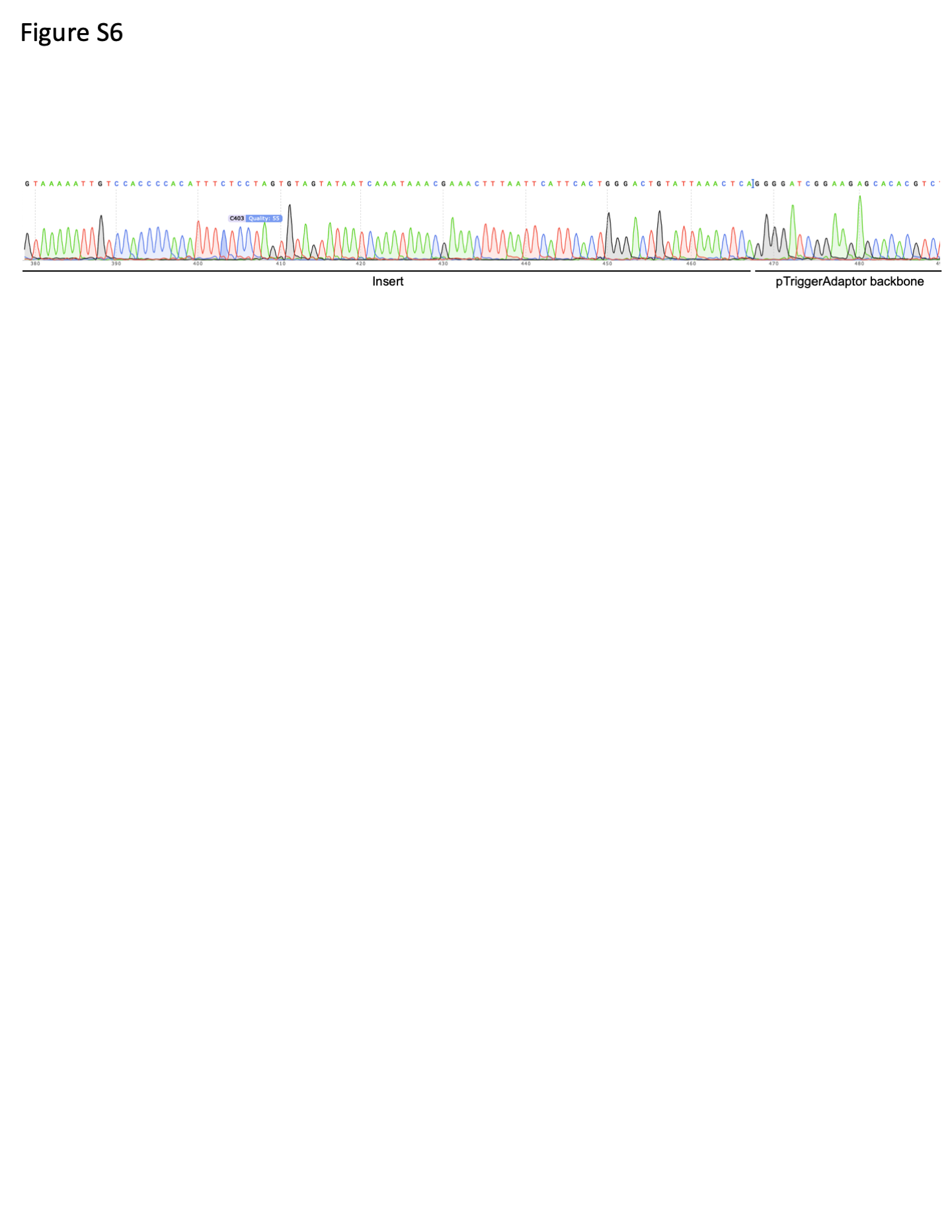

### Figure S7

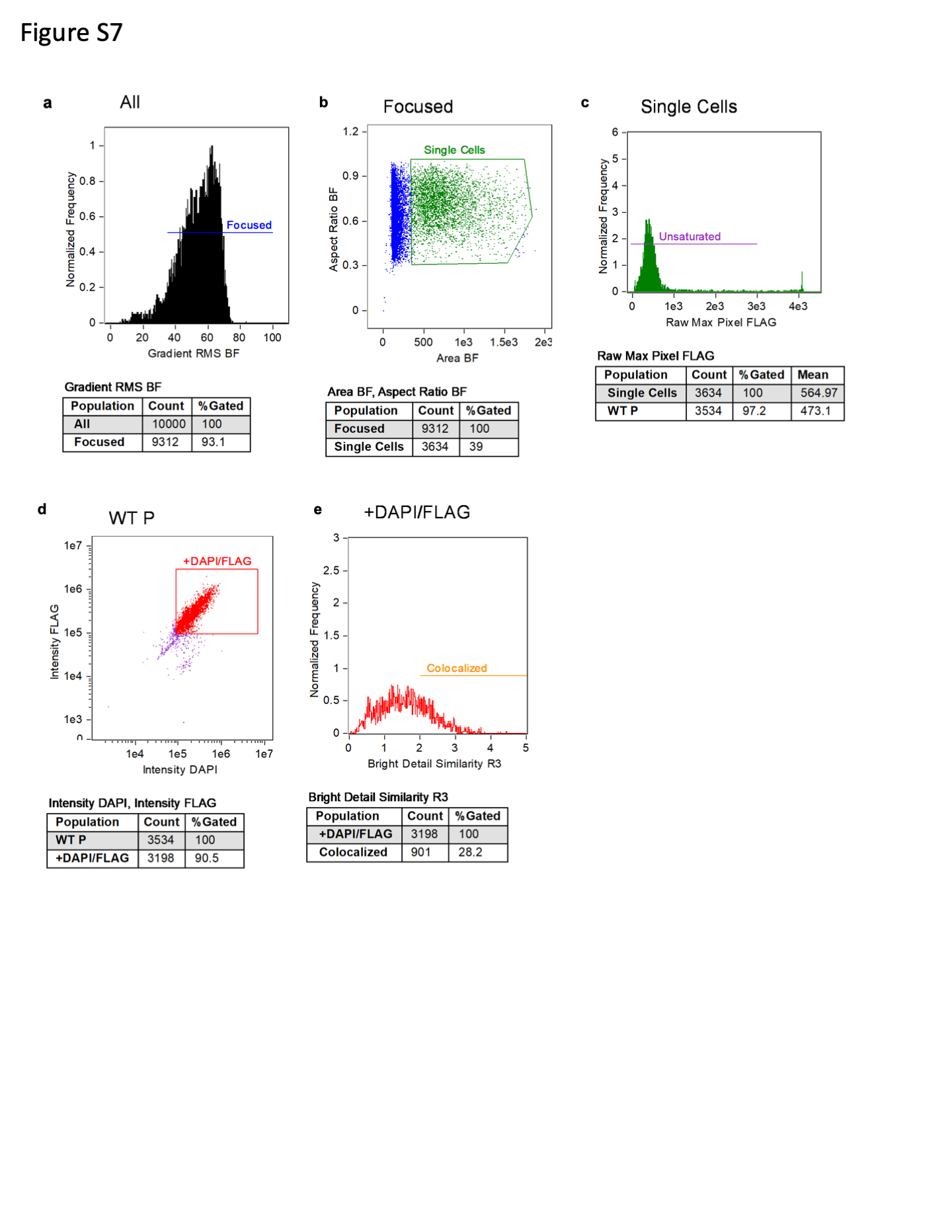

### Figure S8

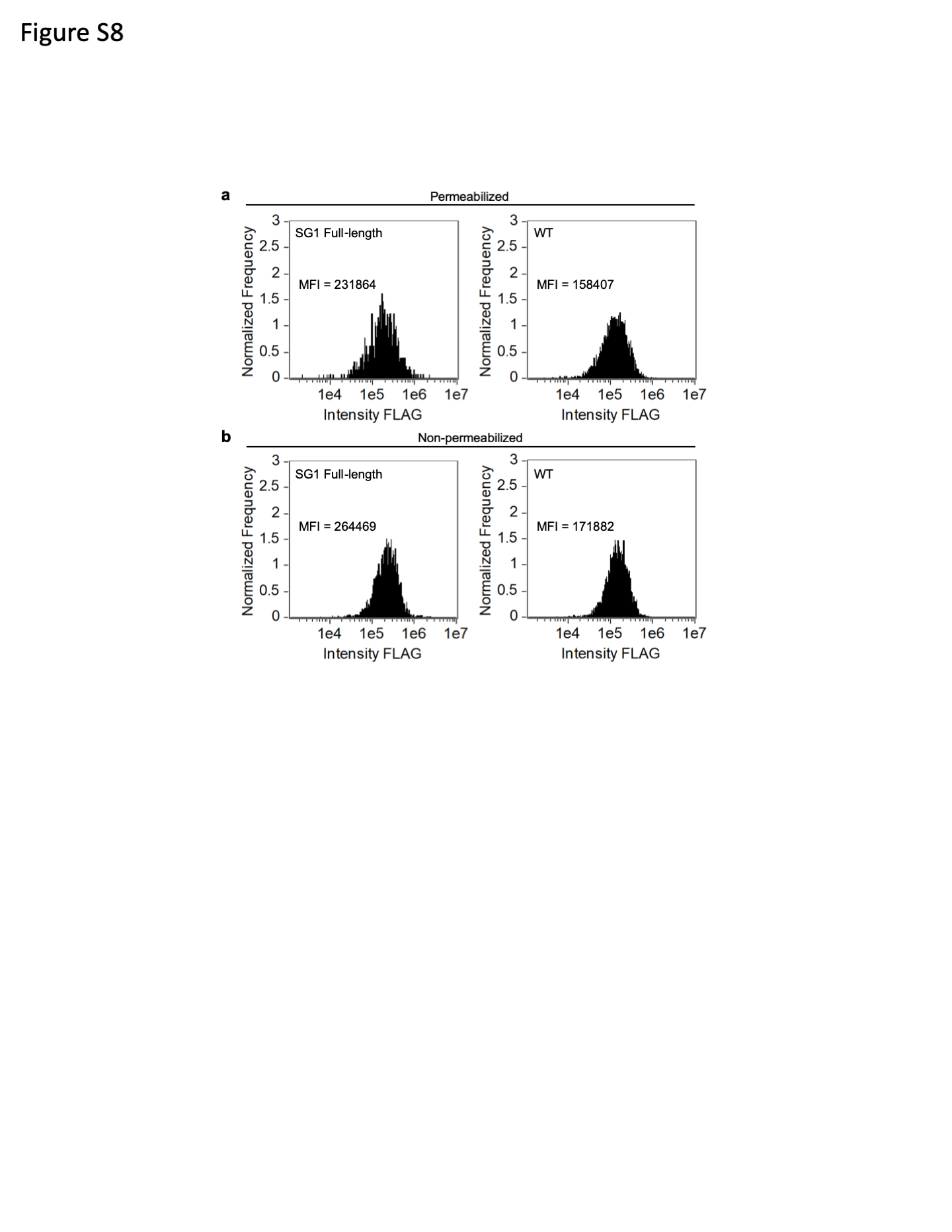

### Figure S9

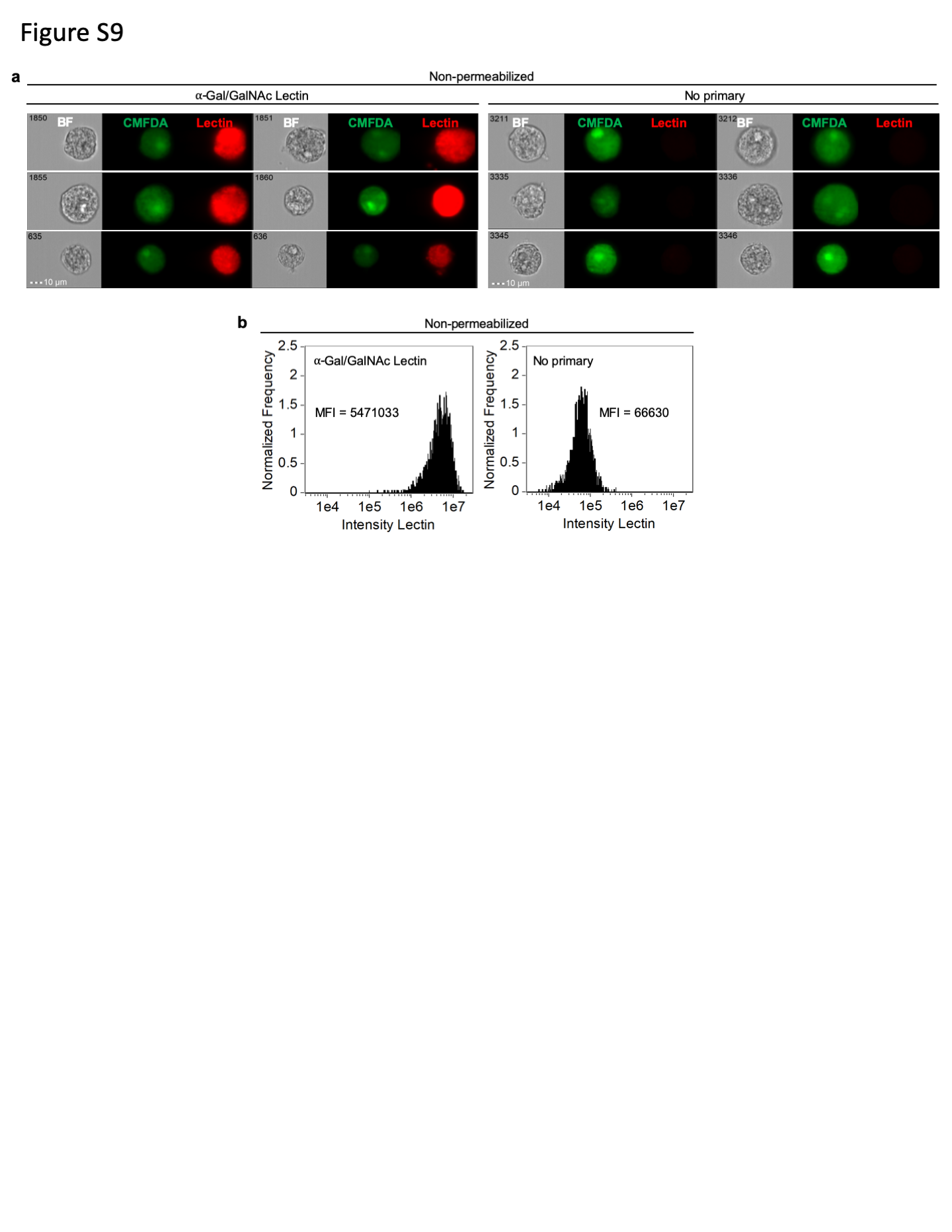

### Figure S10

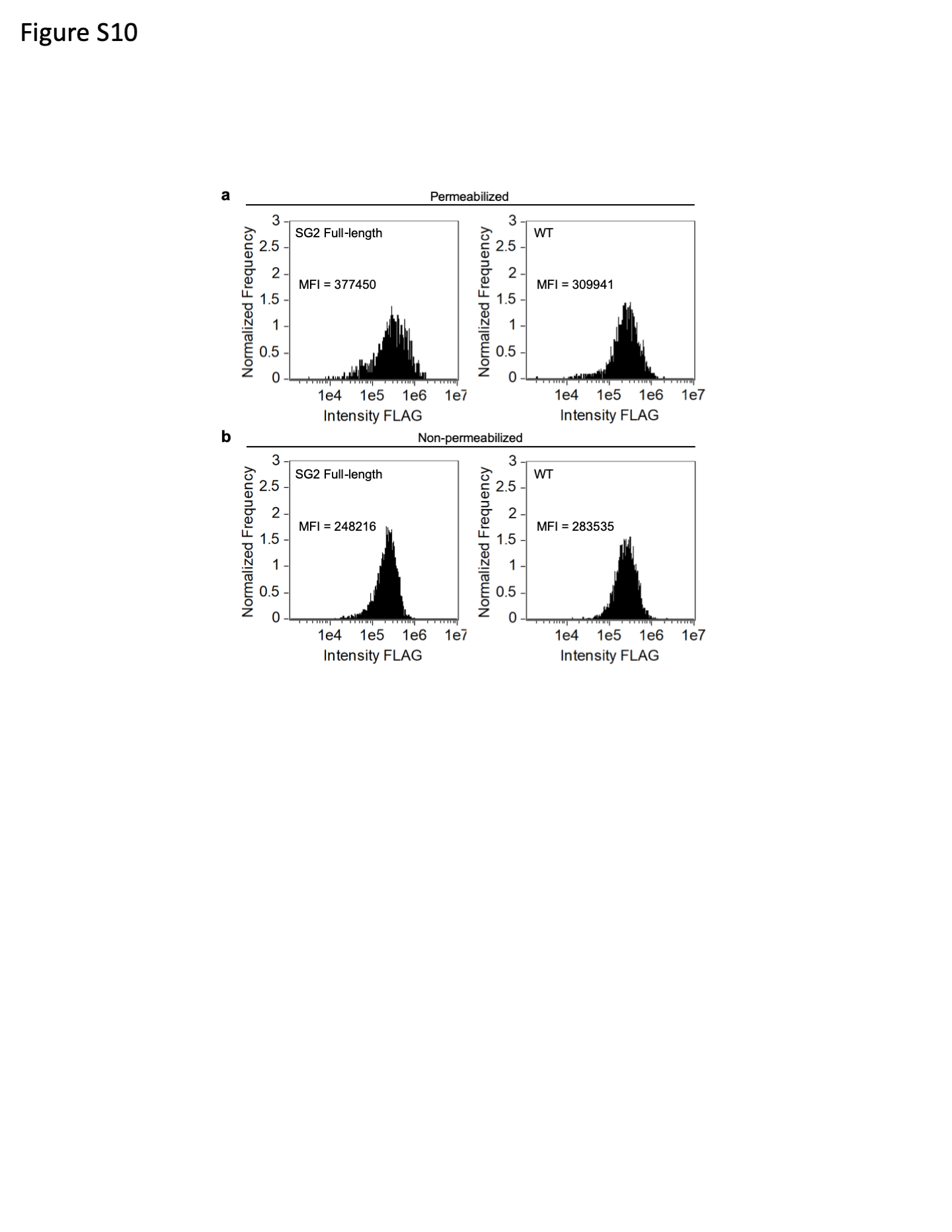

### Figure S11

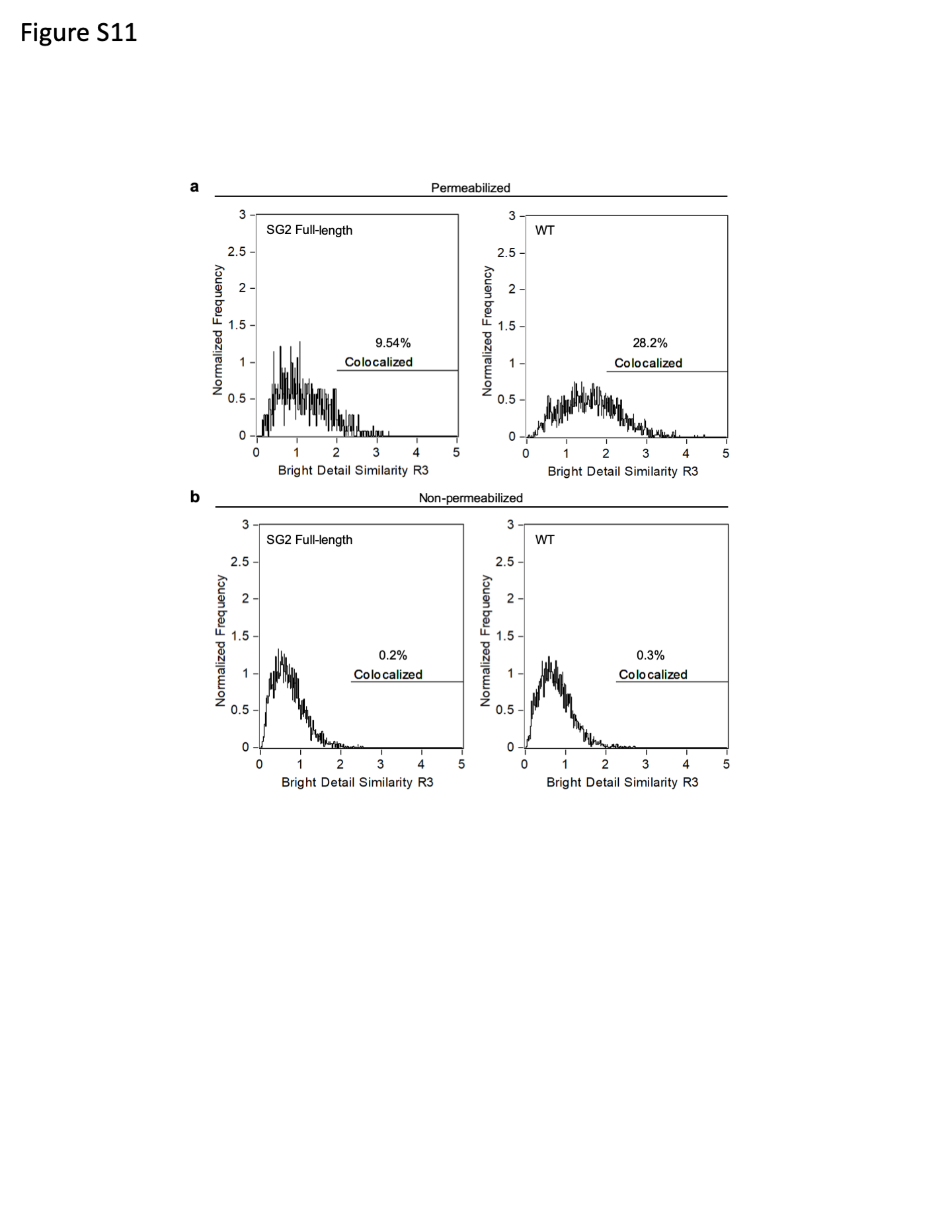

### Figure S12

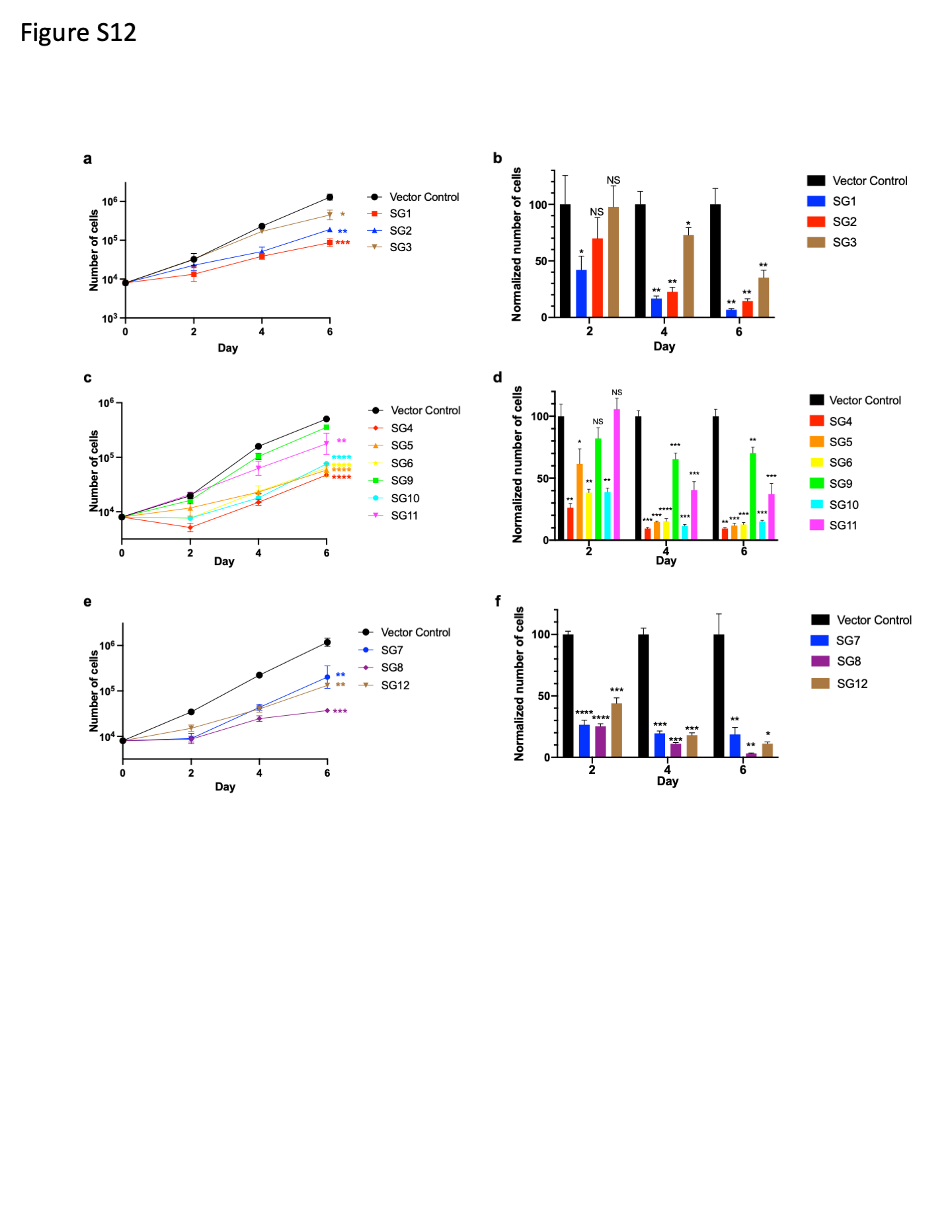

### Table S1

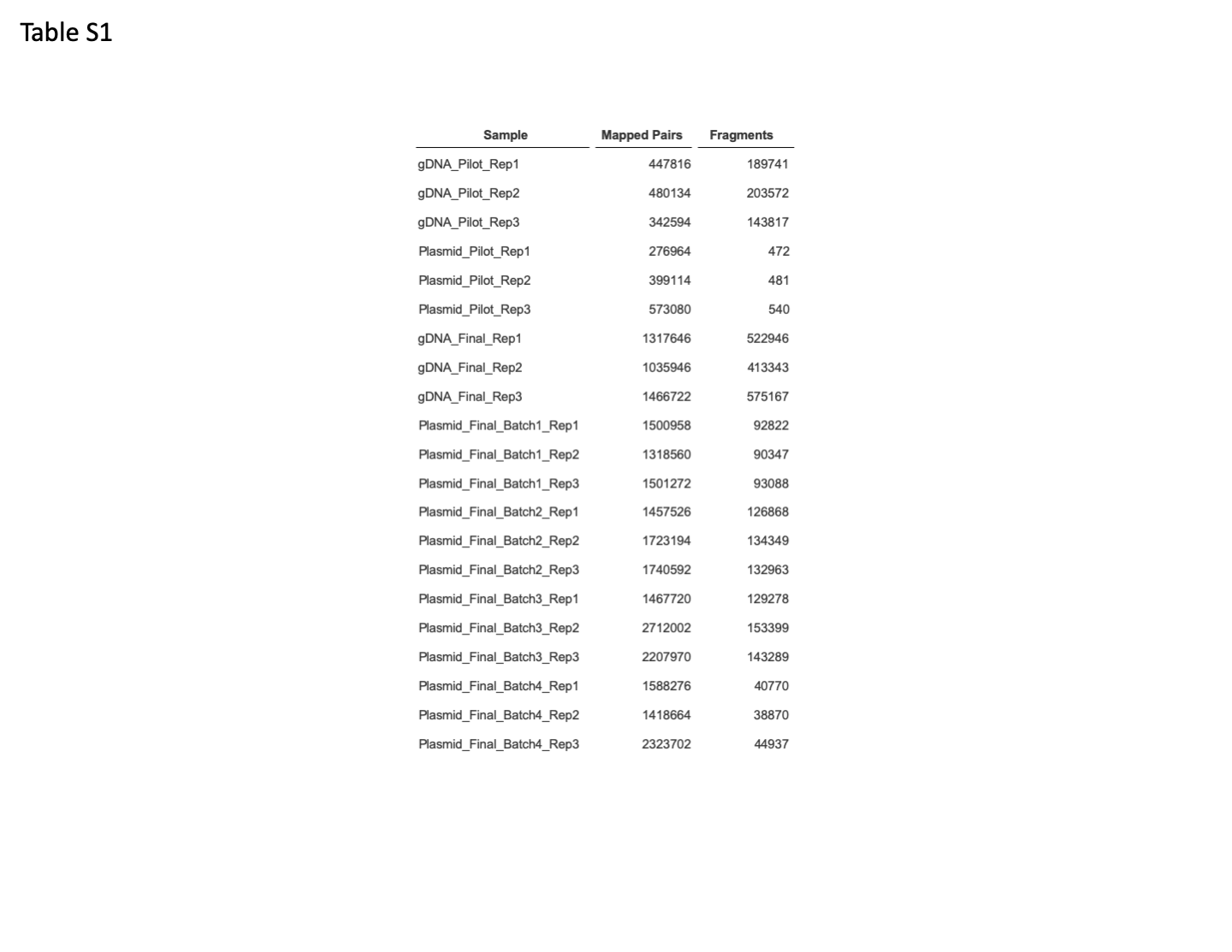

### Table S2

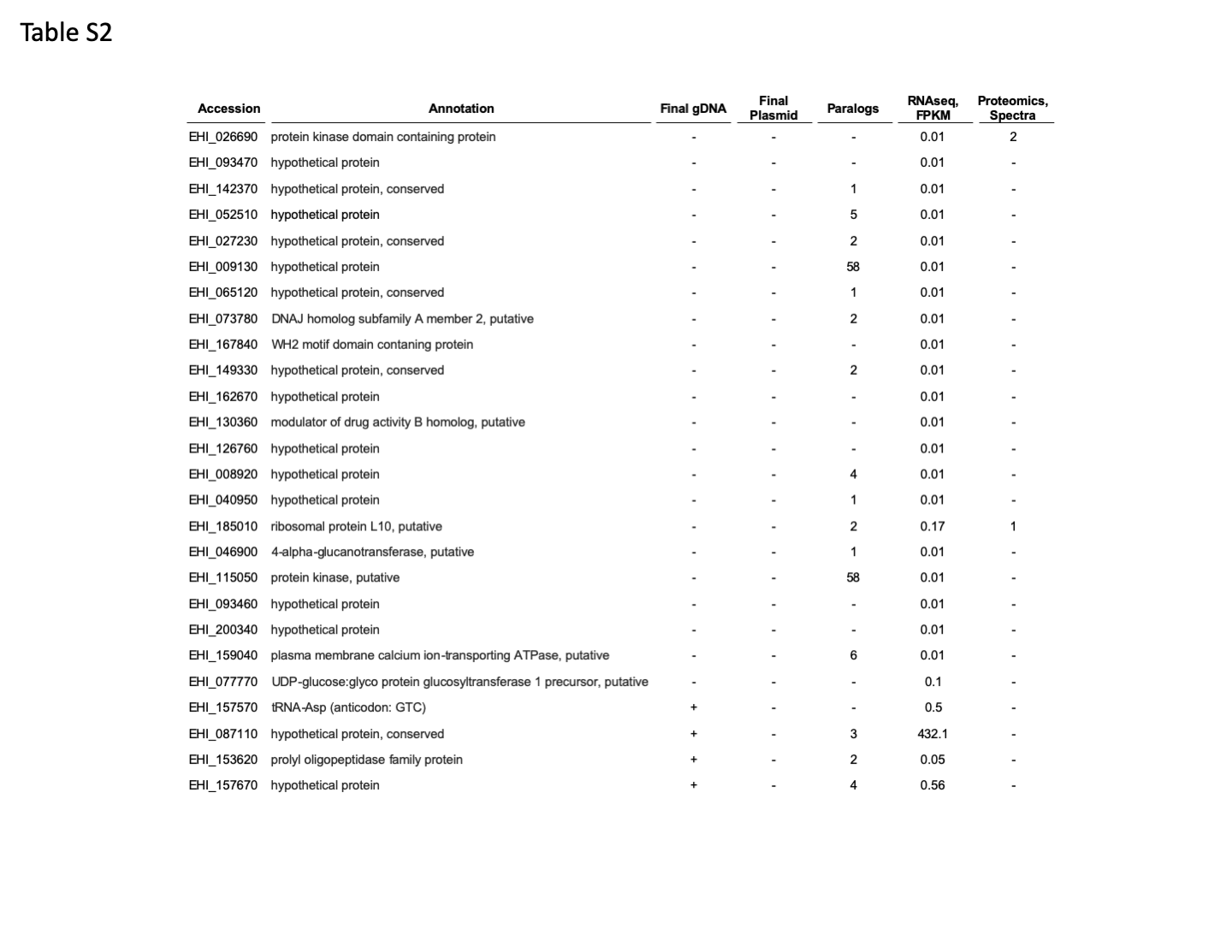

### Table S3

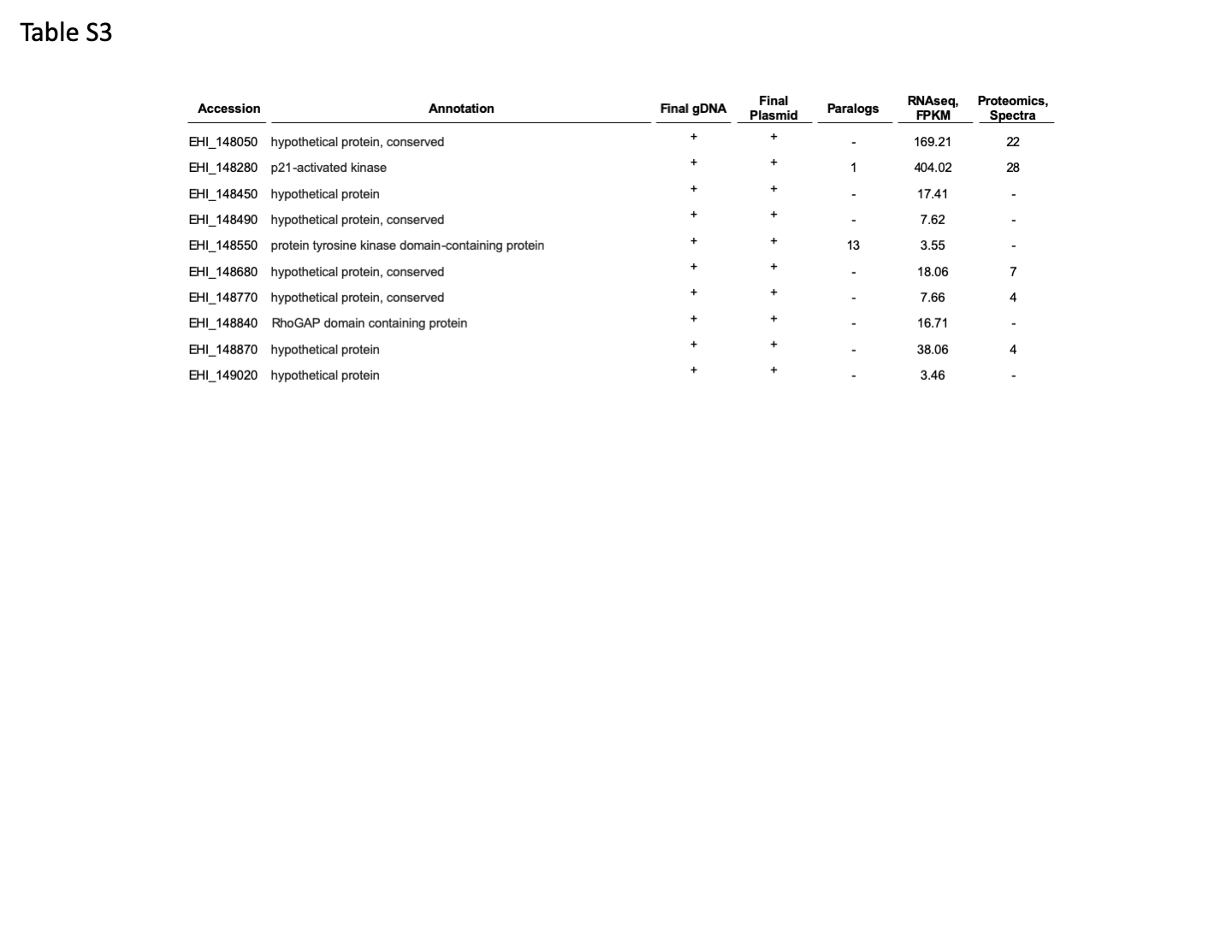

### Table S4

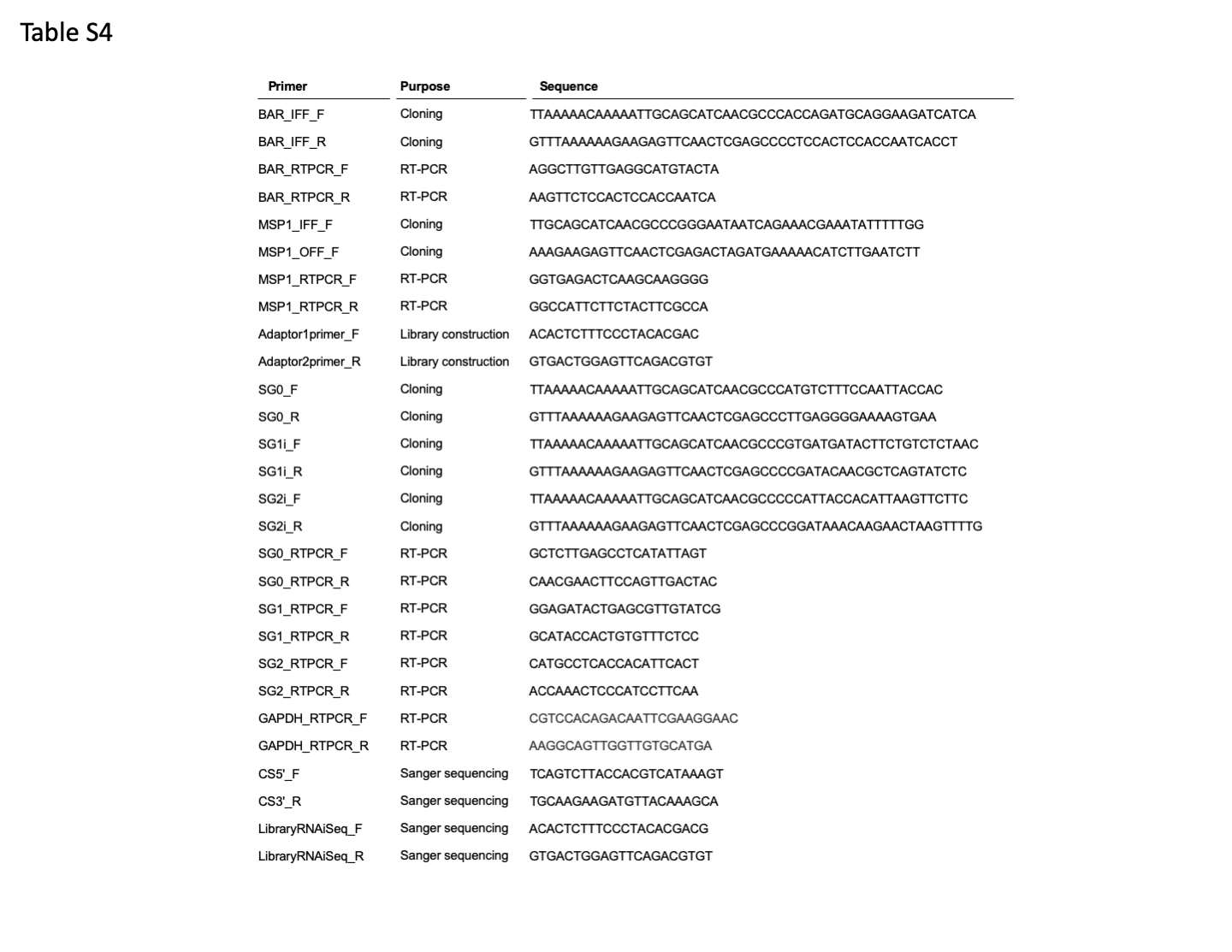
